# *Nematostella vectensis* exemplifies the exceptional expansion and diversity of opsins in the eyeless Hexacorallia

**DOI:** 10.1101/2023.05.17.541201

**Authors:** Kyle J McCulloch, Leslie S Babonis, Alicia Liu, Christina M Daly, Mark Q Martindale, Kristen M Koenig

## Abstract

**Background:** Opsins are the primary proteins responsible for light detection in animals. Cnidarians (jellyfish, sea anemones, corals) have diverse visual systems that have evolved in parallel with bilaterians (squid, flies, fish) for hundreds of millions of years. Medusozoans (e.g. jellyfish, hydroids) have evolved eyes multiple times, each time independently incorporating distinct opsin orthologs. Anthozoans (e.g. corals, sea anemones,) have diverse light-mediated behaviors and, despite being eyeless, exhibit more extensive opsin duplications than medusozoans. To better understand the evolution of photosensitivity in animals without eyes we increased anthozoan representation in the phylogeny of animal opsins and investigated the large but poorly characterized opsin family in the sea anemone *Nematostella vectensis*.

**Results:** We analyzed genomic and transcriptomic data from 16 species of cnidarians to generate a large opsin phylogeny (708 sequences) with the largest sampling of anthozoan sequences to date. We identified 29 opsins from *N. vectensis* (*NvOpsins*) with high confidence, using transcriptomic and genomic datasets. We found that lineage-specific opsin duplications are common across Cnidaria, with anthozoan lineages exhibiting among the highest numbers of opsins in animals. To establish putative photosensory function of *NvOpsins*, we identified canonically conserved protein domains and amino acid sequences essential for opsin function in other animal species. We show high sequence diversity among *NvOpsins* at sites important for photoreception and transduction, suggesting potentially diverse functions. We further examined the spatiotemporal expression of *NvOpsins* and found both dynamic expression of opsins during embryonic development and sexually dimorphic opsin expression in adults.

**Conclusions:** These data show that lineage-specific duplication and divergence has led to expansive diversity of opsins in eyeless cnidarians, suggesting opsins from these animals may exhibit novel biochemical functions. The variable expression patterns of opsins in *N. vectensis* suggest opsin gene duplications allowed for a radiation of unique sensory cell types with tissue-and stage-specific functions. This diffuse network of distinct sensory cell types could be an adaptive solution for varied sensory tasks experienced in distinct life history stages in Anthozoans.

## Background

Opsins are a family of G protein-coupled receptors (GPCRs) associated with animal visual systems [1–2]. Previous phylogenetic analysis has shown that bilaterian and cnidarian opsin groups are interspersed and sister to each other, suggesting the major branches of this protein family already diversified in the common ancestor of these lineages, with 3-4 opsins thought to be already present [1,3–6]. In all characterized opsins, the protein binds to a vitamin-A derived chromophore, forming a rhodopsin complex, responsible for absorbing light [2–7]. The transduction of light by rhodopsin into a cellular signal occurs via a G protein signaling pathway known as the phototransduction cascade, leading to downstream ion exchange across the membrane and dictating the cell’s physiological response [2]. Opsins from distinct subclades bind to specific G protein alpha subunits (Gt, Gq, Gs, etc.), which in turn signal via distinct phototransduction cascades [4,8,9]. Eye-associated opsins and their phototransduction cascades are well-studied in vertebrates and insects [2,10–12]. The emergence of expanded sequencing resources across the <u>animal tree of life</u> has led to the discovery of new types of opsins. This has changed our understanding of opsin evolution and has modified historic interpretations of opsin function [13–16]. Although eye-related opsins have received most attention, the importance of non-ocular opsin gene expansions, from which eye-associated opsins have repeatedly evolved, is relatively unexplored [5–17].

The best studied opsins are those expressed in vertebrate or fly eyes. Ciliary associated (c-) opsins were once thought to be exclusive to the vertebrate eye, and microvillar associated rhabdomeric (r-) opsins were thought to be exclusive to protostome eyes [8,18–20]. C-and r-opsins are now known to be expressed in both deuterostome and protostome lineages [14,20–23] and new opsin clades have been found throughout Bilateria (xenopsins and tetraopsins, [24–29]). Cnidarian opsins are grouped in three clades: cnidopsins, which are found throughout Cnidaria, and two **a**nthozoan-**s**pecific **o**psin clades (ASO-I and ASO-II) [6–29]. Cnidopsins are sister to the bilaterian-specific xenopsins. The positions of ASO-I and ASO-II are not consistently supported in published available opsin phylogenies [1,6,29–31]. ASO-I has been found either to be sister to r-opsins or sister to all animal opsins [1,6,29–31]. ASO-II is considered sister to ciliary opsins, although this is not well supported [1,6,30,31]. However, ASO-II shares intronic structure with ciliary opsins, which supports this placement [29]. Together these data suggest at least 3 major opsin clades existed in the ancestor of both Bilateria and Cnidaria [1,4,6,15,32]. Evidence from both eyed and eyeless cnidarians shows that cnidarian opsins function as photoreceptors using canonical phototransduction cascades [3,31,33–46]. Both sequence homology and physiological evidence suggest cnidarian opsin biochemistry and function is similar to bilaterian opsins [47].

Among anthozoan opsins, gene family expansions have been observed, yet the phylogenetic history and potential adaptive consequences remain unknown [6–29]. Although anthozoans lack eyes, large numbers of opsins have been reported in corals and sea anemones [6]. Recent next-generation sequencing in Cnidaria has improved phylogenetic sampling and clarified relationships among all animal opsins. However, cnidarian opsins have undergone large gene family expansions and transcriptomic studies have failed to consistently identify the numbers of opsin genes in each lineage, especially within anthozoans [5,6,47]. This variability is likely due to similar sequences of recent duplicates, spatiotemporal restriction or low expression. Without complete genomes and manual annotation, the extent of cnidarian opsin expansions and potential functional diversification remains obscured.

The starlet sea anemone, *Nematostella vectensis,* is an eyeless anthozoan and an ideal model for interrogating the evolutionary history and potential functional diversity of opsins. *N. vectensis* exhibits multiple light-mediated behaviors including spawning, larval swimming, and circadian locomotor activity [48–49]. Previous work has shown *N. vectensis,* like other cnidarians, has many opsins; however, estimates of the precise number of opsins encoded in the genome of this animal have ranged from 30 to 52 [5,6,47]. Furthermore, key requisites for inferring distinct functions of each *NvOpsin*, such as genomic architecture, complete sequence, and whether opsin genes are silent or expressed, have not been analyzed. With the release of two chromosome-quality genomes, we are now able to corroborate transcriptomic evidence with genomic loci in this highly duplicated gene family [50–51]. In this study we reduce the most recent estimate of NvOpsins from 52 [6] to 29, closer to previous estimates [47]. Using evidence from genomic, transcriptomic, phylogenetic, and *in situ* mRNA expression analyses, we characterize a diverse repertoire of opsins from *N. vectensis* and suggest unique populations of opsin-expressing sensory cells have diverse roles across life stages in this animal.

## Results

### Phylogenetic identification and genomic architecture of the 29 Opsins in N. vectensis

Using multiple genomic and transcriptomic sources of evidence, we were able to identify 159 additional opsin sequences from 15 recently reported anthozoan transcriptomic and genomic datasets, with each species having between 4 and 22 paralogs (Fig 1A). We report the chromosomal location of all previously reported *NvOpsin* gene models while removing erroneous predictions and adding new loci, identifying a total of 29 distinct genes (Fig 1B). To facilitate comparisons across previously published investigations of opsin diversification, we have provided a table of *N. vectensis* opsin IDs from previous studies with new genomic identification and an updated naming convention (Additional File 3: Table S2). We made a maximum-likelihood gene tree for the opsin protein family, adding *N. vectensis* sequences and additional new anthozoan data to published cnidarian opsin alignments (Additional files 1, 2). We used this tree to investigate the orthology of NvOpsins (Figs. 1-3). ASO-I opsins are highly supported (99.9 /100 SH-aLRT/UFboot support) within the opsin clade, sister to other opsins (Fig1C, for full tree support values see Additional File 2). [5,6,29]. Cnidopsins and xenopsins form a monophyletic group (88.6/90 SH-aLRT/UFboot support) and the ASO-II group is found sister to c-opsins, however this relationship is not highly supported (Fig. 1C). Our tree shows all NvOpsins fall into ASO-I, ASO-II, and cnidopsin subclades (Fig. 1C).

**Figure 1.**
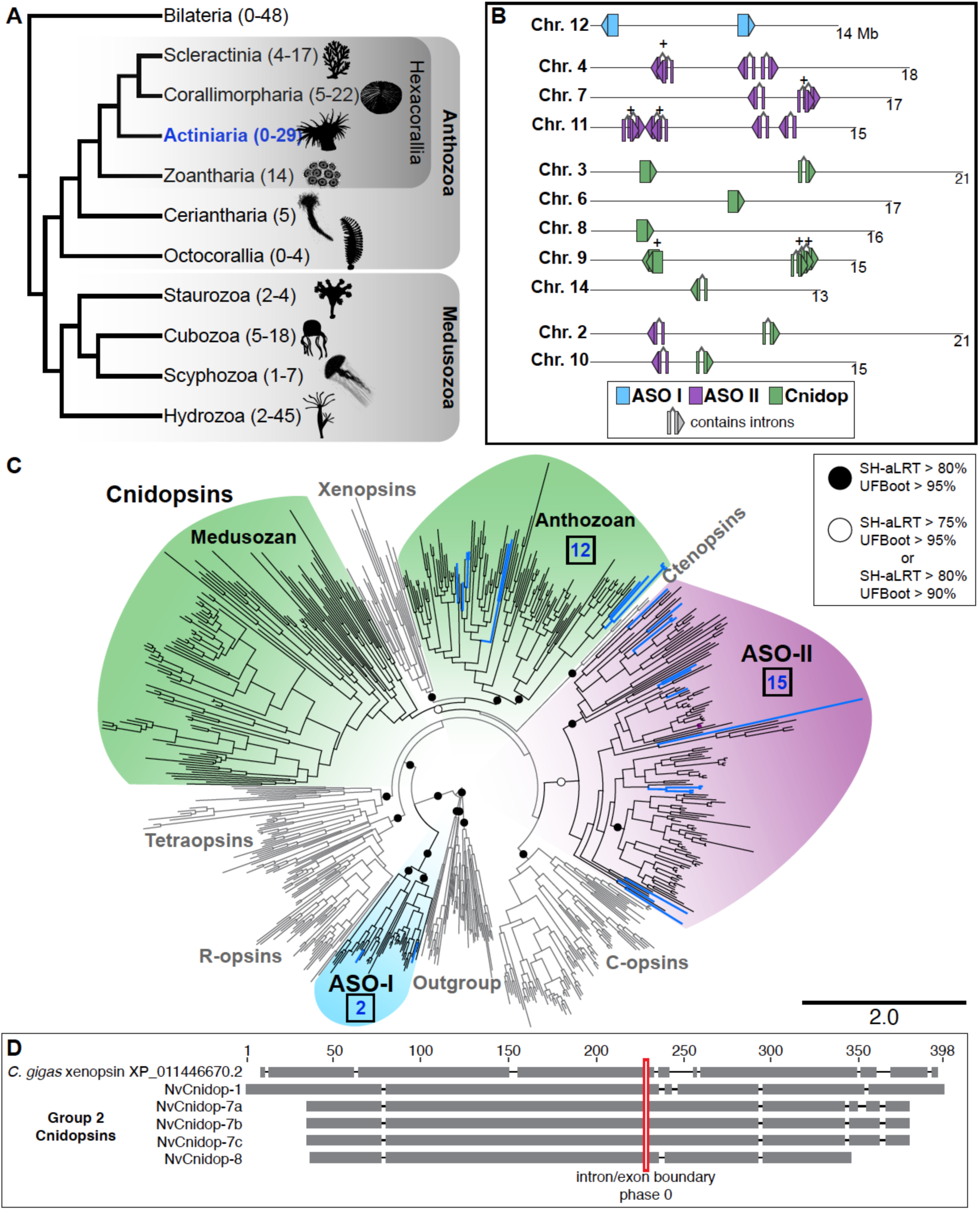
Phylogenetic placement and genomic architecture of the 29 *N. vectensis* opsins. **A)** Simplified cnidarian phylogeny adapted from [87]. The range of known opsins found in a single species in each group is indicated on the phylogeny. *N. vectensis* is in Actiniaria, Hexacorallia, in the Anthozoa (in blue) and has the most opsins so far identified of any anthozoan (N = 29). (**B,C**), light blue corresponds to ASO-I, purple to ASO-II, and green to cnidopsins. **B)** *N. vectensis* chromosomes with opsin loci are shown. Nearly all *N. vectensis* opsins segregate on chromosomes by clade. Numbers below each chromosome are length in megabases, (+) indicates recent tandem duplicates with highly similar sequences. Arrowheads indicate direction the gene is found in the genome. **C)** Maximum likelihood tree of 708 opsins, with major animal opsin clades labeled. Cnidarian-specific clades are colored and in bold. IQtree branch support is defined by ultrafast bootstraps (Ufboot) and likelihood ratio test (SH-aLRT). The number of *N. vectensis* opsins in each clade is listed (blue numbers). **D)** Conservation of intron structure in the cnidops/xenops clade. A representative xenopsin from the oyster *Crasssostrea gigas* shares an intron/exon boundary with all other xenopsins investigated [29] and several *N. vectensis* cnidopsins. Red box shows intron/exon boundary mapped on to amino acid alignment. Gray bars represent aligned sequence, black lines are gaps in the alignment.

To further investigate possible mechanisms of opsin diversification, we assessed genomic structure and conservation of intron-exon boundaries of opsin genes in the *N. vectensis* genome (Fig. 1D, Additional File 2). Investigating chromosomal locations of all *NvOpsin* loci, we found that chromosomes containing multiple opsins largely contain genes from the same subclade and not from other subclades (Fig. 1B). We further examined intron/exon structure in *NvOpsins* and found that both *NvASO-I* genes and 5 of the 12 *NvCnidopsins* lack introns, a hallmark of duplication by retrotransposition (Fig 1B, Additional File 3: Table S2) [29,38,40]. By contrast, all *NvASO-II* genes and the remaining 7 *NvCnidopsins* have introns [50]. Together, these results suggest tandem duplication and retrotransposition have been major drivers of opsin expansion in *N. vectensis*.

To further examine the evolutionary relationships within the xenopsins/cnidopsins clade, we examined intronic structure in *N. vectensis* and representative bilaterians. Cnidopsins together with xenopsins form a single clade suggesting one ortholog was present in the cnidarian/bilaterian ancestor and that each lineage (anthozoa, medusozoa, and bilateria) experienced extensive lineage-specific duplication of this ancestral gene. We compared NvCnidopsin intron/exon boundaries to bovine c-opsin and a *Crassostrea gigas* xenopsin. Anthozoan cnidopsins are split into two subclades (see below), and all *NvCnidopsins* in Subclade 2 contain introns (Additional File 3: Table S2). None of these cnidopsins share intron/exon boundaries with c-opsin, but all Group 2 *NvCnidopsins* share one intron/exon boundary position and phase with the oyster xenopsin (Fig. 1E). This conserved intron-exon structure further supports a common ancestor of all cnidopsins and xenopsins and suggests Subgroup 2 *NvCnidopsins* are structurally like the ancestral cnidopsin.

### Extensive duplication/divergence of opsins is a common feature of Hexacorallia

The addition of more anthozoan sequences in our phylogeny reveals opsin gene family expansions are common in Hexacorallia (Fig. 2). In general, all anthozoans have two ASO-I opsin paralogs that group into two monophyletic subclades, suggesting that two ASO-I opsins were present in the common ancestor of this clade. Very few lineages exhibit more than two ASO-I paralogs although one or both duplicates are absent in several lineages, and both *NvASOI* duplicates are intronless, suggesting gene loss, rather than duplication, shaped the evolution of ASO-I opsins (Fig. 2A). In contrast, ASO-II opsin paralogs are expanded, and divergence within and between lineages is high (see branch lengths, Fig. 1C). ASO-II duplications already occurred in the lineage leading to Hexacorallia (Fig 2B), while additional recent duplications greatly expanded ASO-II numbers within Hexacorallia (Fig. 2A,B). No ASO-II members have been identified in any Octocorallia species (soft corals, sea pens), suggesting a loss in this lineage. The presence of two ASO-II sequences from ceriantharians (tube anemones) suggests a minimum of two ASO-II opsins were present in the last common ancestor of Hexacorallia. Within the ASO-II clade, previously identified ASO-II 2.1 and 2.2 subgroups [6] are well supported. Previously identified Subgroup 1 is less well supported, and this large group shows four duplication events occurred in the lineage leading to Scleractinia and Corallimorpharia. Among sea anemone opsins, parallel duplications have occurred between the major anemone subfamilies (Anenthemonae – to which *N. vectensis, Scolanthus, and Edwardsiella* belong, and Enthemonae to which all other included species belong) (Fig. 2A, Fig. 3B). Subgroup 2.1 has fewer duplications in hexacorallian lineages than Subgroup 1. Sister to Subgroup 2.1 is a newly identified group (Subgroup 3) including the highly divergent NvASOII-4 sequence (Fig. 3B, 4B), orthologs of which were identified in Corallimorpharia and Scleractinia but not found in any other sea anemone. Previously Subgroup 2.2 was thought only to contain sea anemone sequences [6]. The addition of new data identifies cerianthid, zoanthid, and corallimorpharian sequences, suggesting this subgroup of opsins arose in the last common ancestor of Hexacorallia (Fig. 3B).

**Figure 2.**
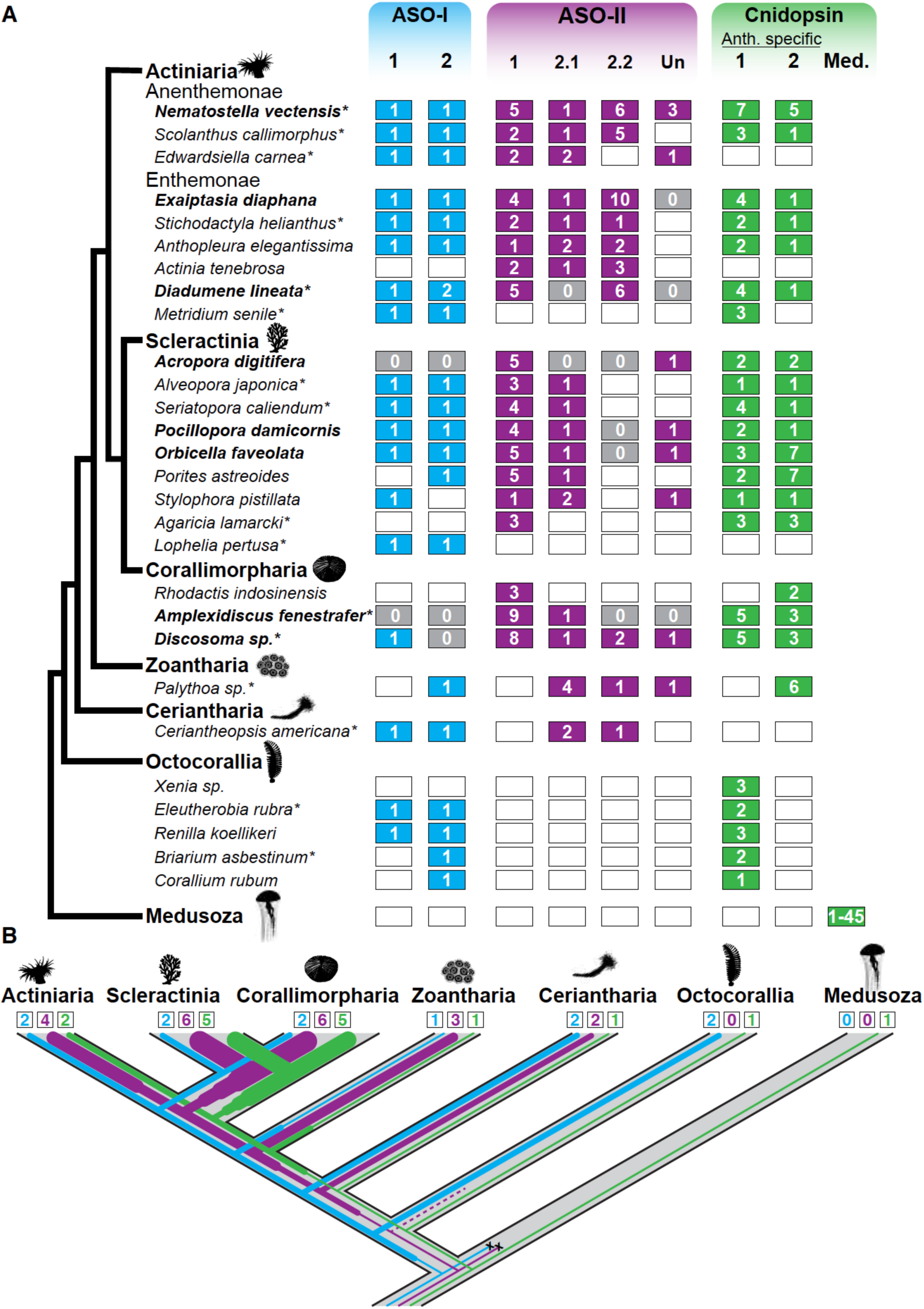
Current and ancestral anthozoan-specific opsin duplications. **A)** Anthozoan species with opsin data are listed. Each column is the subclade number within the three major anthozoan opsin clades. “Un” is ASO-II opsins that are unspecified, or not found in previously identified clades. The number of opsins for each species and opsin subclade is listed in the boxes. Gray boxes with 0 signify genomic evidence of no opsins, while white boxes signify no opsin identified from transcriptomic evidence. Species names in bold have genomic evidence available, asterisks are for species with newly reported opsins in this study. **B)** Anthozoan lineage tree, with numbers of opsins represented by line thickness and estimated by parsimony based on opsin numbers and phylogenetic positions (Fig. 3). Numbers in boxes represent the estimated number of opsins present in the last common ancestor of each anthozoan lineage. An increase in thickness of lines signifies an opsin duplication while an x signifies loss of the gene. The dashed line signifies uncertainty due to limited sampling in Octocorallia.

**Figure 3.**
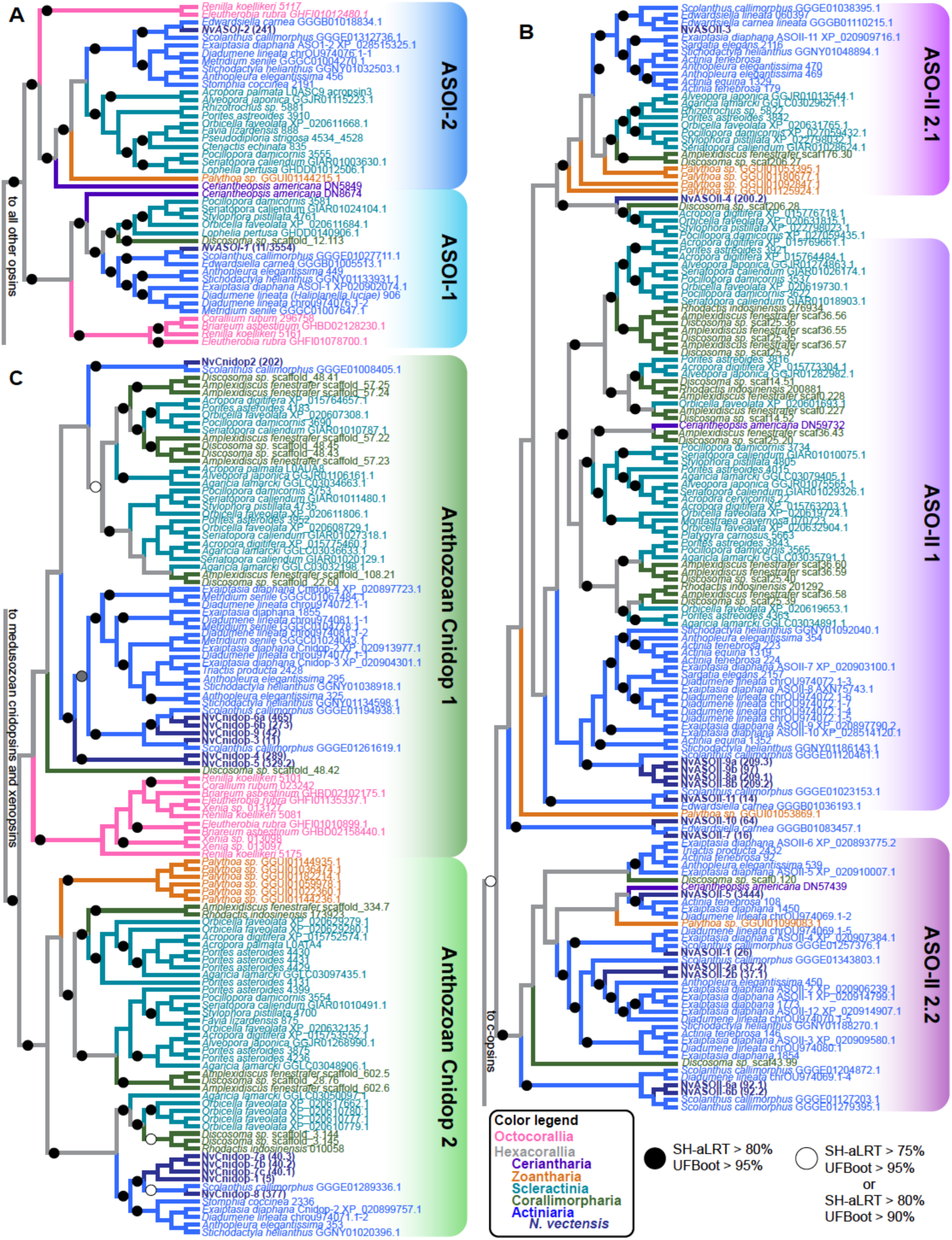
Anthozoan-specific opsin evolutionary patterns of duplication and loss. Trees are zoomed subsets of the maximum likelihood tree (Figure 1C) for each anthozoan opsin group. Species names and branches are color coded according to lineage. Support for branches is denoted with a black circle or a white circle. **A)** ASO-I group opsins are split into two main subclades, with most anthozoans having an opsin duplicate in each subclade **B)** The ASO-II group is sister to c-opsins and comprised exclusively of opsins from hexacorals. Previously identified ASO-II Groups 2.1 and 2.2 [6] are well-supported but ASO-II Group 1 is not. **C)** Anthozoan cnidopsins form a single well-supported clade within the larger cnidopsin/xenopsin clade. Within Anthozoan cnidopsins there are two sister clades, one of which is well-supported while the other is not. For clarity, branch lengths are transformed, and branch support is not shown for branches leading to the two shallow-most nodes on these trees (For full tree topology and support values see Fig. 1C, Additional File 2).

Within the cnidopsin clade, we identify one well-supported subclade containing only Hexacorallia and another clade containing all anthozoan lineages (Fig. 2, Fig. 3C). This suggests one cnidopsin ortholog was present in the common ancestor of anthozoans and a duplication event occurred in the common ancestor of Hexacorallia. Like the ASO-II group, multiple lineage and species-specific duplications have expanded cnidopsin numbers in anthozoans. Cnidopsin Subgroup 1 contains all the octocorallian sequences in a single well supported clade, with at least one octocorallian-specific duplication event or multiple species-specific events. No cerianthid sequences were identified as cnidopsins, though one is likely placed improperly in the bilaterian-specific xenopsins, and likely to be the cerianthid cnidopsin ortholog (Additional File 2). The ancestor of Scleractinia and Corallimorpharia duplicated their Group 1 and Group 2 opsins at least once, followed later by multiple species-specific duplications (*A. digitifera,* six opsins, *Discosoma,* eight opsins). In *N. vectensis*, lineage-specific expansions led to 12 total cnidopsins, in parallel to multiple independent duplications in the other sea anemone group, Enthemonae. Together our phylogeny reveals extensive duplication of opsins throughout the diversification of anthozoans contributed to the expansive repertoire of opsins in *N. vectensis* and other cnidarians.

### NvOpsins are intact GPCRs with a potential diversity of functions

In addition to phylogenetic placement, we assessed whether the 29 NvOpsins were capable of functioning as GPCRs and photopigments by analysis of conserved functional residues. The most basic criteria for any opsin to function as a photopigment are the presence of seven transmembrane domains, which form a chromophore binding pocket, and the presence of a lysine at position 296 (bovine rhodopsin numbering), which binds to the chromophore (Fig. 4A) [2]. The presence of a pair of cysteines to stabilize the opsin structure via a disulfide bridge is additionally required (Fig. 4A) [52]. A negatively charged residue known as the counterion is thought to be required to stabilize the interaction with the chromophore, but the specific site and residue of this counterion can vary across opsins clades [2]. Beyond these basic functional features, conserved sites on the cytoplasmic tail/loops bind to specific G protein alpha subunits, which are required for activating the phototransduction cascade [53–54]. Opsins within a clade are usually specialized in binding one or limited number of G protein alpha subunit subtypes (Gt, Gq, Gs, etc.) [8]. The few bilaterian and jellyfish opsins that have been characterized in detail have identified specific functionally relevant amino acid sites and identities required for G protein signaling [41–42] (Fig. 4A).

**Figure 4.**
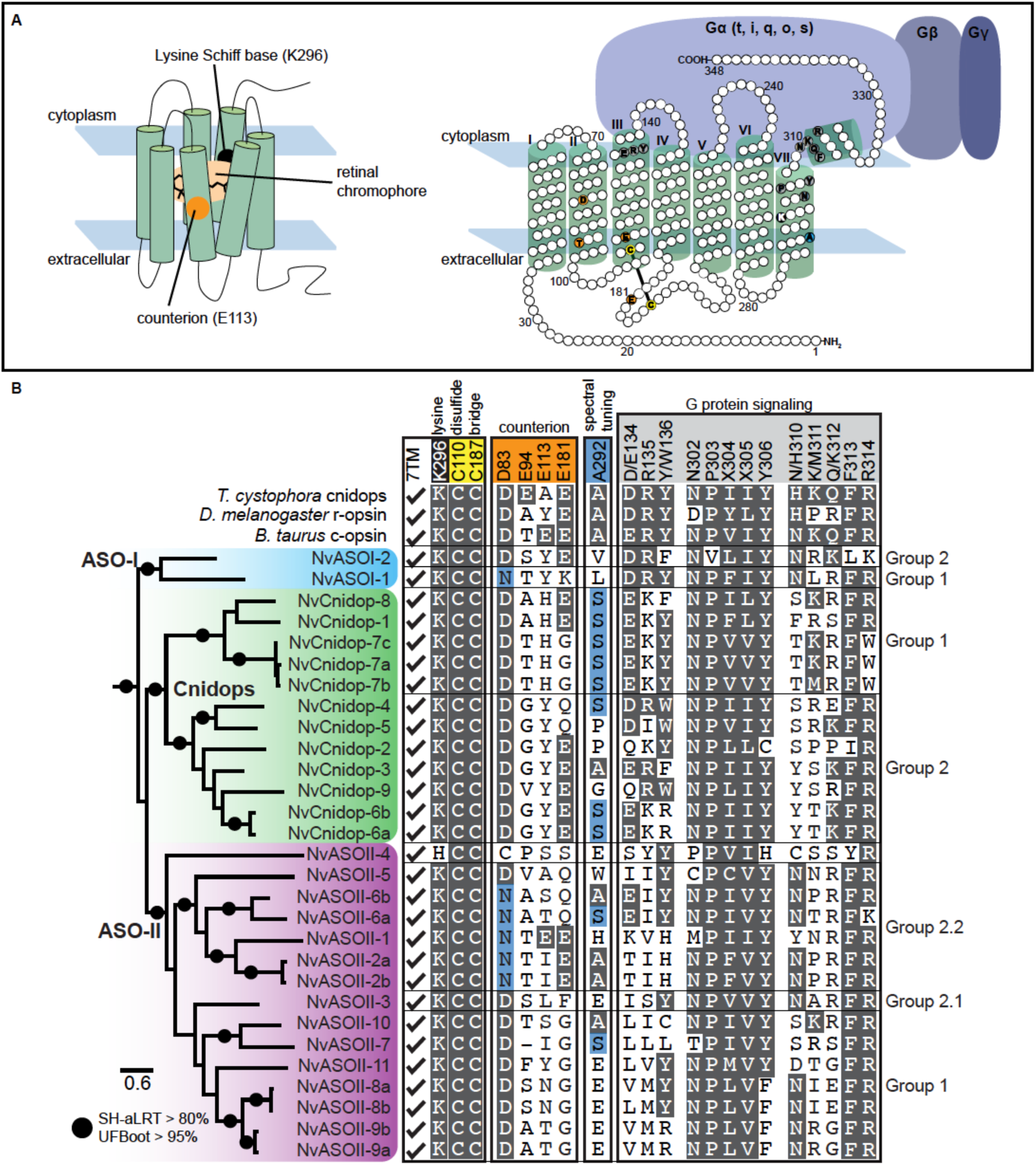
*N. vectensis* opsins have high levels of sequence diversity at canonically conserved functional sites. **A)** Left, cartoon bovine rhodopsin structure showing 7 transmembrane domains surrounding the vitamin A derived chromophore and the highly conserved lysine at position 296 (black) required for chromophore binding. The glutamic acid counterion (orange) is conserved among vertebrate c-opsins and important for chromophore binding. Right, cartoon diagram of bovine rhodopsin sequence with select functional sites color coded by functional category, matching the numbered sites in (B). Cartoon G protein subunits are shown bound to rhodopsin. The specific G protein alpha subunit can vary (letters in parentheses) depending on opsin sequence, with functional implications for type of signaling cascade activated. **B)** Left, maximum likelihood phylogeny of *N. vectensis* opsins, with IQtree2 support. Right, select *N. vectensis* opsin amino acids are aligned with bovine rhodopsin (c-opsin), *D. melanogaster* r-opsin, and *T. cystophora* cnidopsin. Canonically conserved functional residues and positions follow bovine rhodopsin numbering and correspond to (A). From left to right, the first box contains structural features minimally required for function in all opsins. 7TM indicates protein sequence has seven transmembrane domains. Black is the conserved Lys(K)296; yellow shows conserved cysteine residues that form a stabilizing disulfide bridge. Second box lists known counterions from bilaterian and box jelly opsins (orange). Third box shows a conserved spectral tuning site across opsin clades (blue). A second spectral tuning site is also found at counterion site Asp(D)83. Substitutions known to cause a blue shift in both sites are shaded blue. Fourth box contains conserved sites known to be important for G-protein signaling (light gray). NvOpsin residues that are conserved at the known functional sites are shaded in dark gray. Rightmost, NvOpsins are labeled by subclade within each major anthozoan opsin clade.

All NvOpsins have the canonical seven transmembrane domains and all but one have the conserved lysine, Lys-296 (Fig. 4B). Interestingly, NvASOII-4 has a Lys296His substitution at this site, suggesting a non-photoreceptive function (Fig. 4B). Previously identified counterion sites included Glu-113 in vertebrate ciliary opsins and Glu-181 in invertebrate Go-coupled opsins and Gq coupled r-opsins [8–55]. There is some evidence that Asp-83 is the counterion site for Gq-coupled melanopsin, the r-opsin found in vertebrates, and Glu-94 is the counterion in cubozoan Gs-coupled cnidopsin [8–41]. No NvOpsin has Glu-94 (cubozoan) or similar negatively charged amino acid at this position, however some have either Glu-181 or Glu-113 and NvASOII-1 have both (Fig. 4B). All NvCnidopsins, most NvASO-II, and one NvASO-I sequence has Asp-83, while other NvASO-II and NvASO-I sequences has an Asn substitution at the Asp-83 site. Only NvASOII-4 has neither aspartic acid nor asparagine in this position, instead having Cys-83 (Fig. 4B). Together sequence analysis suggests NvOpsin paralogs tend to have bilaterian-like counterion residues while the Glu-94 counterion appears to be an evolutionary novelty in box jellies.

Spectral tuning – the shift in the spectrum of wavelengths an opsin absorbs – can be mediated by amino acid substitutions that affect the interaction with the chromophore [56]. Two substitutions, Asp83Asn and Ala292Ser, result in opsin absorbance shifts from green toward blue, and have evolved independently in multiple vertebrates [57–59] and insects [60–61]. We find that in many *N. vectensis* sequences, one or both substitutions are also present, and evolved independently within *N. vectensis* opsins multiple times (Fig. 4B). This suggests potential blue shifts in wavelength absorption compared to related opsins, although the absorbance spectra of these opsins remain to be measured. The functional relevance of other amino acids at either of these sites is unclear. The conservation of these sites in opsins as distantly related as c-and r-opsins suggests their presence is also involved in spectral tuning of *N. vectensis* opsins, but further functional evidence is needed for confirmation.

We also investigated conservation of sites known to be important in G-protein signaling. In the rhodopsin-like GPCR family, the conserved tripeptide Asp/Glu-Arg-Tyr/Trp at sites 134-136 is known to be important for receptor activation, located after the third transmembrane domain in the second intracellular loop [62–64]. Only NvASOI-1 and NvCnidop-4 have all three of any of these residues. Most NvCnidopsins have a similarly positively charged lysine in place of arginine at site 135, which may be a conservative substitution. We also searched for three known motifs that directly interact with the G protein alpha subunit and are important for G protein signaling activation [65]. Most *N. vectensis* opsins have NPXXY at these sites, but some of the NvASO-II sequences are divergent at this highly conserved motif. Another known functional motif is the tripeptide at sites 310-312 followed by a conserved FR sequence (313-314). The canonical c-opsin and r-opsin tripeptide is NKQ and HMK respectively, while the functional box jelly photopigment, Tcop13, has HKQ at this site [40]. In the box jelly opsin*, in vitro* substitution with other tripeptides did not result in inactivation of Gs signaling, but instead altered the dynamics of light response in cell lines [40]. The FR motif is conserved in NvOpsins, but none of the known tripeptide motifs are found in any NvOpsin. Given the *T. cystophora* evidence, these data may suggest functional diversity in binding dynamics of G proteins across NvOpsins.

### Opsin expression is dynamic throughout N. vectensis tissues during development

To investigate which tissues may play a role in light-sensitive behaviors, we analyzed spatiotemporal expression patterns of opsin expression in *N. vectensis* using a combination of RNAseq and *in situ* hybridization. The lifecycle of *N. vectensis* proceeds from fertilization to blastula stage by 12 hours and gastrula stage by 24 hours. By 48 hours a swimming planula larva stage develops [66]. This stage is characterized by an aboral ciliated sensory organ known as the apical organ, which faces forward when swimming. By 10 days post fertilization mesenteries and four tentacles form and elongate and the planula metamorphoses into a primary polyp.

We generated a high-quality *de novo* assembled transcriptome from four developmental stages (blastula, gastrula, mid-planula, and primary polyp) as well as male and female adults. Out of 221,245,806 total reads, 96.29% were properly paired and used for downstream applications with a BUSCO completeness of 97.9%. Replicate libraries of the same stage were highly concordant with one another (Additional file 4: Fig. S1). The RNA-seq time course analysis revealed that *NvASOI-2* is highly expressed at blastula stage relative to all other opsins (Fig. 5A, top scale). To better visualize the diversity of expression of the remaining opsins we show the rest of the expression data scaled without this highly abundant transcript (Fig. 5A, bottom scale). The time course analysis allows for qualitative comparison across developmental stages for each opsin transcript. Many NvOpsins peak in expression at different stages, including blastula (*NvASOII-8a, -9a*), gastrula (*NvASOII-9b*), planula (*NvASOII-3; NvCnidop-1, -3, -9*), and primary polyp (*NvASOII-2a, -2b -3, -6a; NvCnidop-2b, -4, -6b*) (Fig. 5A). We also compared our data with a published developmental time course (NvERTx) for all opsins found in that dataset [67], which generally agreed with our expression levels and provided additional time resolution at early stages (Additional file 4: Fig S2). Light is known to induce spawning in *N. vectensis* and other cnidarians, so we wanted to know whether some opsins were specific to adults or specific to sex. Our comparison between adult males and females showed several opsins that were expressed in both adult sexes as well as six opsins that were differentially expressed by sex. *NvASOI-2, NvCnidop-2*, and *NvCnidop-8* were upregulated in females, while *NvASOII-6b* and *NvCnidop-4* were more highly expressed in males (Fig. 5B). Together the RNA-seq data suggest dynamic expression of distinct NvOpsins throughout development and across sexes.

**Figure 5.**
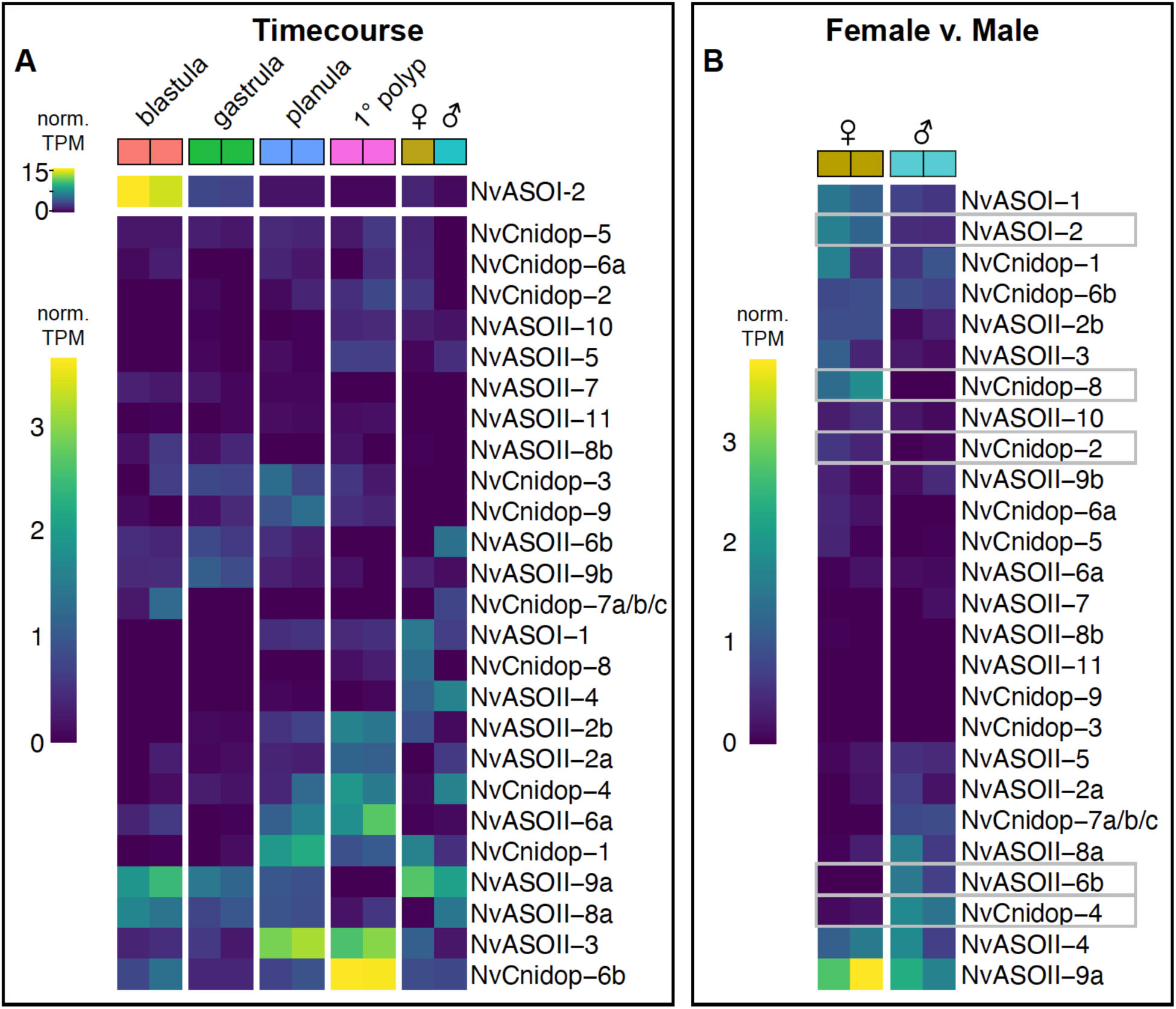
NvOpsins are expressed dynamically throughout development and between sexes. **A)** Expression levels from time course analysis are plotted for opsins across all stages and both sexes. Each box is from a single replicate library. For visual clarity, highly expressing *NvASOI-2* is shown using a separate scale. Heatmap illustrates variable opsin expression across development in all opsin transcripts. *NvCnidop7a/b/c* loci are highly similar in coding sequence, such that transcripts are not distinguished between the three sequences. *NvASOII-1* had no high-identity transcript in our transcriptome or in NvERTx. **B)** A separate analysis comparing male and female adult opsin expression is visualized in a heatmap. The gray boxes indicate opsins that were significantly differentially expressed between sexes, with a significant q-value of < 0.05.

To corroborate our RNA-seq analysis and better understand tissue specificity, we performed *in situ* hybridization for select opsin genes at peak expression levels (Fig. 6). Our expression studies show that tissue-and stage-specific expression patterns do not correlate with phylogenetic signal. Opsins of distinct clades are found in both germ layers and sometimes have overlapping expression patterns. Opsins from each major clade were expressed in both embryo and polyp stages. (Figs. 5-6).

**Figure 6.**
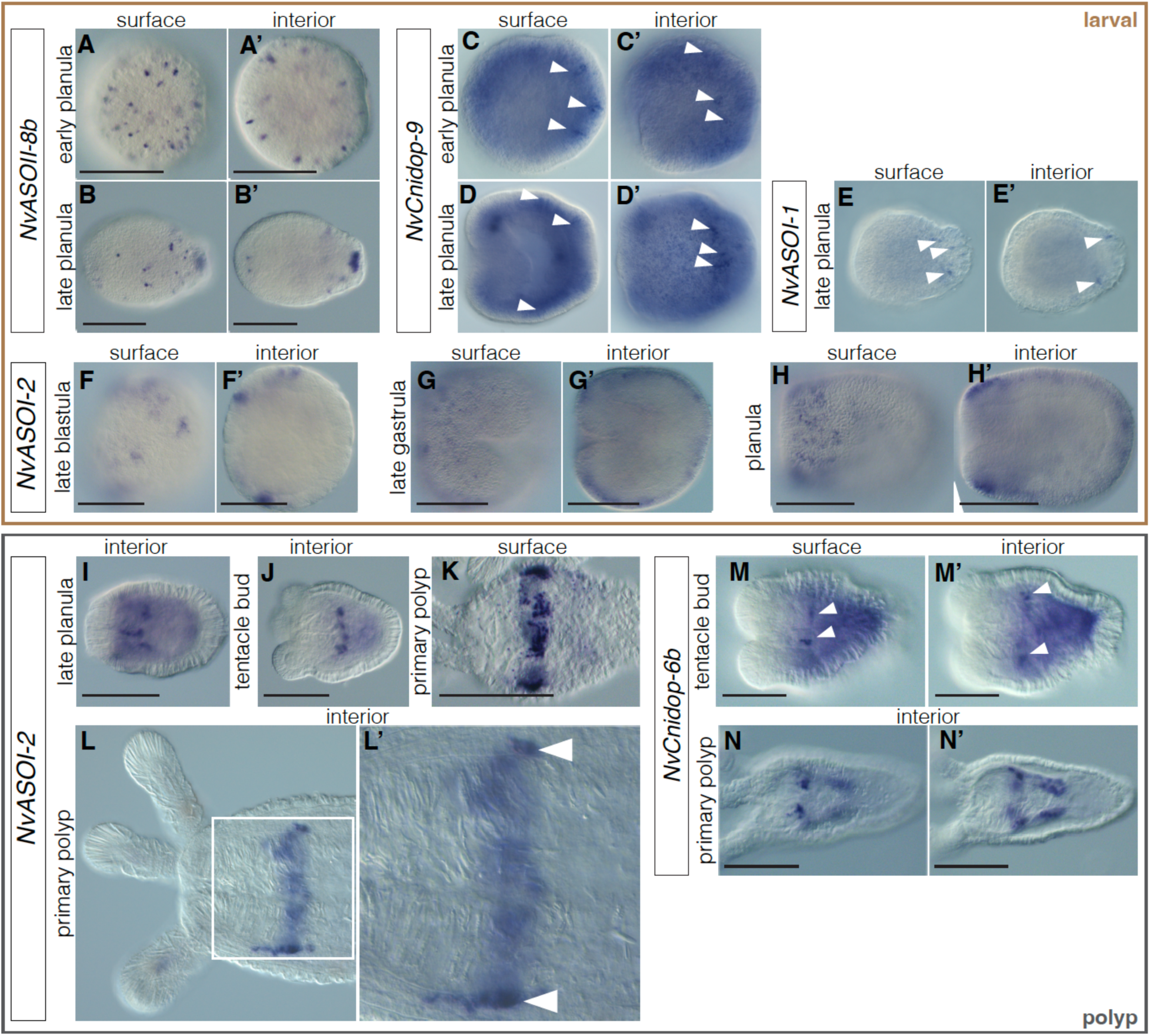
NvOpsin spatial expression patterns suggest a diversity of functions throughout development. **A-B’)** *NvASOII-8b* is only expressed at the swimming planula stage. Expression is ectodermal, in individual cells scattered throughout the ectoderm in early planula. In later stage planula (B’) the expression is also concentrated at the aboral end in the sensory apical organ. **C-D’)** *NvCnidop-8* is expressed in a subset of aboral ectodermal cells at planula stage (arrowheads). **E-E’)** *NvASOI-1* is also expressed in a subset of cells in the aboral ectoderm. **F-H’)** *NvASOI-2* is expressed in the ectoderm at early stages. Starting at blastula stage (F) through gastrula (G), expression is patchy in clusters of cells which tend to be concentrated more orally later in development (H). By late planula/tentacle bud stage expression has concentrated in an endomesodermal ring at the aboral end of the pharynx, where it can be seen into the primary polyp stage. **I-L’)**). **M-N’)** *NvCnidop-6b* begins to express at late planula/tentacle bud stage in the pharyngeal endomesoderm, and by polyp stage is expressed in specific cells throughout the mesenteries. Scale bar 100 um; in all images oral is left, aboral is right.

In embryonic and larval stages, we observed opsin expression in both ectoderm and endomesoderm. *NvASOII-8b* is expressed by gastrula stage in scattered cells throughout the ectoderm (Fig. 6A,A’) and by late planula, there is also expression in the sensory apical organ (Fig. 6B,B’). Both *NvCnidop-9* and *NvASOI-1* are expressed in the ectoderm at planula stages (Fig. 6C-E’) and are expressed in scattered cells of the aboral ectoderm but are not expressed in the apical organ. *NvASOI-2,* the opsin with the highest expression in the RNA-seq dataset, is detected in scattered cells at late blastula stage and its expression expands into the ectoderm in larval stages (Fig. 6F-H’). Expression is concentrated orally as development continues, eventually forming a ring around the oral region in the late planula stages (Fig. 6H-H’).

*NvASOI-2,* which is expressed in ectoderm in larval stages, is found in endomesodermal tissue surrounding the pharynx at the time of tentacle bud formation and into the primary polyp stage (Fig. 6I-L’). *NvCnidop-6b* is also expressed in clusters of cells within the pharyngeal endomesoderm and extends aborally into the mesenteries. (Fig. 6M-N’). Distinct tissue-and stage-specific expression of opsins from distinct subclades suggests different functions throughout development.

## Discussion

We have identified 29 opsins in the genome of *N. vectensis,* the most of any anthozoan so far investigated [6–38]. Maintenance of complete coding sequences, evidence of distinct genomic loci for similar paralogs, and expression patterns lead us to conclude with high confidence these encode *bona fide* opsin proteins (though some may have non-canonical opsin function). Our addition of recent anthozoan phylogenetic data has revealed new patterns of duplication and evolution, both in understudied cnidarian lineages and poorly characterized opsin clades. *N. vectensis* shows large lineage-specific expansions in both cnidopsins and ASO-II opsins. While *N. vectensis* opsins are generally conserved with related bilaterian opsins, significant divergence at canonically conserved “essential” sites suggests high functional diversity. The investigation of anthozoan opsin clades not found in medusozoans potentially leads to novel opsin functional diversity, such as the non-canonical NvASOII-4. This is also reflected in mRNA expression, where multiple opsins are expressed dynamically throughout development, differentially expressed between adult sexes, and spatially restricted to specific cell types and tissues.

Our opsin tree was able to capture more anthozoan-specific opsin evolutionary history than previous work due to increased sampling with recently available sequencing. Our data show expansions in both ASO-II and cnidopsins are concentrated in Hexacorallia and duplications have occurred at both ancient and recent timescales (Fig. 3). A limitation to this and other trees is mRNA sequencing tends to miss opsins expressed at a restricted stage, within a restricted tissue or those that are too similar to be distinguished by assembly algorithms. Our analysis of two chromosome-level genomes was able to add and remove erroneously annotated or previously missed *N. vectensis* opsins. More high-quality long-read genomic sequencing would greatly increase confidence in opsin numbers for many species and help define patterns of opsin duplication and loss in Cnidaria. In addition, variable phylogenetic support for deep opsin relationships between Bilateria and Cnidaria requires methods such as intron analysis to show evidence of shared ancestry. All medusozoan opsins are cnidopsins and nearly all are intronless [40]. The Hydrozoan *Clytia hemispherica* is an exception, with two intron-containing opsins. However, neither of these intron-containing opsins share intron/exon boundaries with any other cnidarian opsin or xenopsin [31–68]). We show for the first time using genomic data from *N. vectensis* that the cnidopsin locus shares a common intron/exon structure with xenopsins in bilaterians. The lack of introns in medusozoan and some *N. vectensis* cnidopsins suggests duplication by retrotransposition may have occurred early in the expansion of cnidopsins or in parallel after Anthozoa and Medusoza split. Furthermore this suggests a medusozoan-specific loss of the original intron-containing genes, while some Anthozoa have maintained this ancestral gene structure shared by xenopsins.

Expression of NvOpsins is dynamic, found across life stages, tissue types, and sexes. *NvASOII-8b,* found only at the swimming stage and in the sensory apical organ, may function in larval light detection and control of swimming. Others, such as *NvCnidop-6b,* are found in the developing mesenteries and upregulated in adult males, relative to females, suggesting a role in reproduction. *NvASOI-2* is highly expressed early and persists through adult stages, its expression domain changes over time, and it is sexually dimorphic in adults. Primordial germ cells develop in the mesenteries near the pharynx, at a similar location to *NvASOI-2* expression [69]. Together this suggests important functional roles throughout development and a possibly distinct adult role in reproduction (Fig. 7). Functional studies in other anthozoans have suggested many light-dependent behaviors may be mediated by specific opsins though none have directly shown this in any anthozoan. In corals, some opsins are expressed at the aboral pole of the swimming larva-similar to *NvASOII-8b* - and were proposed to be involved in light detection while swimming [34]. In coral, some opsins have been proposed to be important for broadcast spawning or algal symbiosis though neither has been shown directly [43–70]. Unlike corals, where husbandry and functional genetics can be challenging, *N. vectensis* is extremely amenable to laboratory study and genetic manipulation. By characterizing the opsins in this animal, future experiments are now able to link behavior and cell physiology to specific opsins.

**Figure 7.**
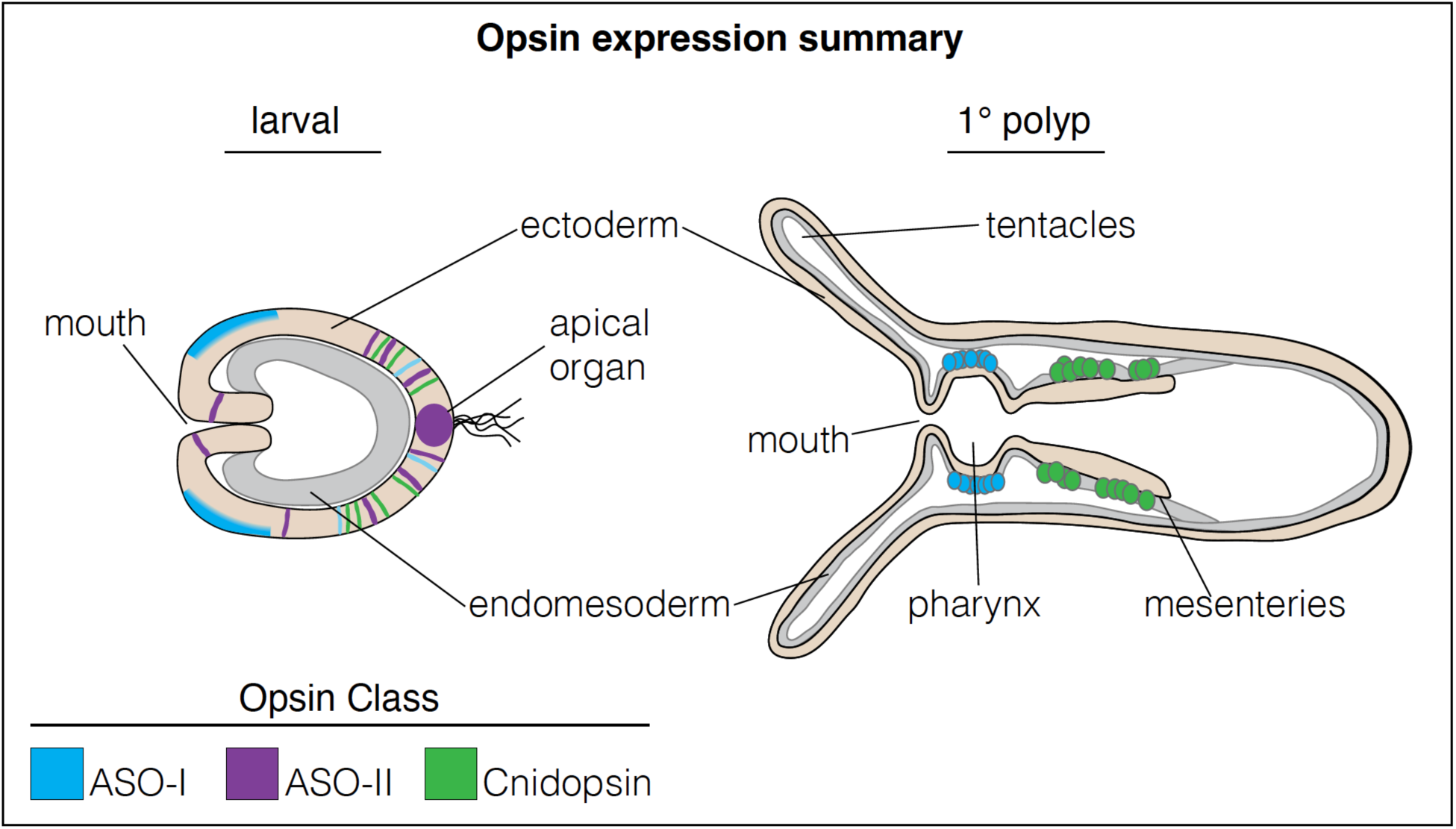
Summary of opsin expression patterns in development. Two summary stages are shown for the primarily early larval expression, and the primarily later primary polyp expression patterns of *N. vectensis* opsins. Opsin expression is color-coded by clade, ectoderm is beige, endomesdoerm is gray. Opsins are expressed throughout developmental stages and tissue types.

Although the divergence from canonically conserved sites suggests new and unidentified functions, we hypothesize that *N. vectensis* has multiple functional opsin photopigments because the animal exhibits several light-mediated behaviors. It is possible that shallow-water species like *N. vectensis* and other anthozoans like reef corals are exposed to more variable light environments, using light sensory information for spawning, substrate settling, defense, and predation. It remains unknown whether opsin gene family expansion in the Hexacoralia is adaptive for these animals, however, our findings suggest a diversity of opsin-expressing cell types involved in multiple light mediated functions in *N. vectensis*.

## Methods

### Animal Care

*N. vectensis* adults were kept in 1/3 concentration artificial sea water in glass dishes on a 12 hour light/dark cycle at 18°C. Animals were fed freshly hatched *Artemia* five times per week. Spawning was induced by placing animals at 24°C overnight with light. Eggs and sperm were collected separately within 2 hours after spawning and fertilized in a new Petri dish and were left to develop at room temperature.

### Opsin Identification and Phylogenetics

Initial *N. vectensis* opsin sequences were collected from three published datasets [5,6,47]. All potential *N. vectensis* sequences were aligned in Geneious Prime v.11.0.12, and identical matches and fragments were discarded. BLAST was used with the final set of opsins as bait to search against the *Nematostella vectensis* Embryogenesis and Regeneration Transcriptomics database (NvERTx) [67], our own reference transcriptome (generated for this study), and two publicly available chromosome level genomes: v2 genome hosted by the Stowers Institute [50–71], and the genome from the Darwin Tree of Life Programme at the Wellcome Sanger Institute (ENA submission accession: ERA9667479) ([51] no associated publication). Hits were added inclusively and combined with a modified alignment including non-redundant sequences from Vöcking et al. 2017, Picciani et al. 2018, and Gornik et al. 2021 opsin phylogenies [5,6,29]. We then added opsin sequences from new anthozoan transcriptome and genome data. We added opsins from 12 species with new transcriptomes, and 3 species from new genomic data (Additional File 3: Table S1). This was done using *N. vectensis* and *Acropora digitifera* sequences from ASOI, ASOII, and Cnidops groups as bait for BLAST searches. The top 100 BLAST hits were initially conservatively kept. All sequences were aligned in Geneious using MAFFT v7.450 with default settings [72]. FastTree was used in Geneious to check the tree, removing unannotated outgroups from the multiple cnidarian and *N. vectensis* BLAST searches. Trimmed (using TrimAl) and untrimmed alignments were generated, but untrimmed sequences generated the best supported phylogeny and are reported here.

Final maximum likelihood gene trees were constructed using IQtree2 with the following command: iqtree2 -s <alignment.phy> -st AA -nt AUTO -v -m TEST -bb 1000 -alrt 1000 [73] on the Minnesota Supercomputing Institute’s (MSI) High Performance Computing system at the University of Minnesota. The LG+G4 model of protein evolution was auto-calculated from this command, and 1000 ultrafast bootstrap (UFboot) replicates and SH-aLRT tests were used for evidence of support. Clades were considered supported only with >80% SH-aLRT/>95% UFboot support. The tree was rooted based on known outgroups from previous phylogenies.

### Genomic and opsin sequence analysis

To identify distinct opsin paralogs, we used BLAST with transcripts as bait to identify genomic loci, resulting in the identification of 29 opsin genes. We manually identified intron/exon boundaries at all cnidopsins with introns by aligning coding sequences to the genome and translated these to identify their location and phase on the protein. We then aligned the cnidopsin amino acid sequences with a xenopsin and bovine c-opsin in Geneious and mapped exon boundaries manually onto the alignment. Opsin genomic coordinates for both new Nvec genomes [51–74], and previous Nvecv1 scaffold coordinates [75] are found in Additional File 3: Table S2. We used InterPro scan in Geneious to predict the seven transmembrane domains. Using bovine rhodopsin as a reference, we aligned our 29 opsins, *Drosophila* r-opsin and *T. cystophora* cnidopsin sequences to identify residues at canonically conserved positions.

A few tandem loci next to *NvASOII-8a, NvASOII-8b, NvASOII-9a, NvASOII9b, NvASOII-7,* and *NvASOII-4* genes were found in the v2 genome [50] whose nucleotide sequences were nearly identical (>97% similarity). We noticed adjacent non-coding intra-and intergenic regions were also near-identical, suggesting these may be haplotype differences and not real paralogs. One of these adjacent pairs also was split by a section of Ns in the genome (Additional File 4: Fig. S3), an indicator of difficulty with assembly in this region. We compared these trouble spots with the Wellcome-Sanger genome [51] and for each, found only one of the two near-identical pairs, further evidence that these copies are a result of assembly error, while other high-similarity but distinct loci were left in our final opsin count.

### RNA-sequencing and analysis

Libraries were generated from two independent spawns each at blastula, gastrula, planula, primary polyp stages, two adult males and two adult females. Adult animals had been kept in normal laboratory conditions at 18°C with a 12 hour light/12 hour dark photoperiod and at least one week since they were last spawned. Live animals were placed directly into Trizol reagent and tissue was homogenized in a microcentrifuge tube with a pestle before performing RNA extraction. Total RNA was extracted from Trizol (Invitrogen), gDNA removed by gDNA cleanup kit (Qiagen), RNA was precipitated using isopropanol and then cleaned up using ethanol precipitation. Libraries were prepared using Kapa Stranded RNA Hyperprep (Illumina), quality control and size selection were performed by the Bauer Core at Harvard University and paired end, 150bp reads were sequenced on an Ilumina NovaSeq.

Raw sequencing libraries were processed for erroneous k-mers and unfixable reads using rCorrector [76]. Adapter sequences were trimmed using Trim Galore (Babraham Bioniformatics, UK), and rRNA reads were mapped to the *N. vectensis* mitochondrial genome and removed using Bowtie2 [77]. All libraries were combined to generate a single *de novo* transcriptome assembly using Trinity v2.12.0 [78]. Transcriptome quality was assessed by checking alignment statistics and quantifying BUSCO completeness [79]. We used EvidentialGene to combine gene and transcript evidence from multiple sources and reduce the number of genes in our transcriptome [80]. We used our own transcripts, and three other assemblies [50,67,81], and the coding sequences from v2 genome [50], as input. We used the EvidentialGene output for our final transcriptome. mRNA quantification was done using Kallisto for each of our 12 libraries using our *de novo* assembled transcriptome [82]. Stages were compared using the time course analysis in Sleuth run in R, and adult libraries were additionally pairwise compared using Sleuth [83]. The data for heatmaps were extracted from the Sleuth object using normalized transcripts per million (TPM) in either the timecourse analysis or male v. female comparison. Heatmaps were made with the heatmap command in gplots package [84] using R v4.2.3 run in RStudio v2022.12.0+353 [85].

### RNA Probe Synthesis

Tissue for probe synthesis was pooled from the same developmental stages as RNA-seq and homogenized in TRIzol (Invitrogen). Total RNA was then extracted with chloroform and cleaned up using a Qiagen RNeasy mini kit. cDNA was made from total RNA using the iScript advanced kit (Bio-Rad). Primers were designed in Geneious (modified Primer3 v2.3.7) (Additional file 3: Table S4). PCR was used to amplify opsin sequences from cDNA and size checked by gel electrophoresis. Properly sized products were ligated to the pGEM T-Easy plasmid (Promega), transformed into chemically competent DH5-alpha *E. coli* cells (New England Biolabs). Plasmid DNA was isolated using the Qiagen miniprep kit and sequenced for confirmation and orientation of insertion into plasmid. Restriction digests linearized the plasmid, were cleaned up using the Qiagen PCR clean up kit, and antisense probes were generated using either Ambion T7 or SP6 megascript RNA polymerases with digoxygenin-labeled (DIG) nucleotides. Probes were cleaned up with Qiagen RNeasy micro kit and eluted in 14 μL. Proper transcription of the probe was checked on a gel. Clean probe was stored at a 50/50 concentration with formamide at -20° C. When ready to use, probe was diluted 1:1000 in hybridization buffer.

### In situ hybridization

*In situ* hybridization was performed according to Wolenski et al. 2013, with slight modification [86]. Before fixation, embryos at blastula and gastrula stages were de-jellied by rocking in 40mg/mL L-cysteine in 1/3x artificial sea water (Instant Ocean) for 10 minutes. Planula and polyp stage animals were immobilized by gently adding 6.5% magnesium chloride in 1/3 seawater. Embryos were fixed in glass vials for 90 seconds in ice cold 0.25% glutaraldehyde/4% paraformaldehyde (PFA) in phosphate buffered saline (PBS). This was removed, 4% PFA was added, and embryos were fixed on a rocker for 1 hour at 4 °C. PFA was washed out with PBS with 0.1% Tween-20 (PTw) and embryos were stored in methanol at -20°C. To begin *in situ* hybridization, embryos were stepped out of methanol and into PTw, then treated with 0.01 mg/mL proteinase K, followed by two glycine washes (2 mg/mL), and two 1% triethanolamine washes in PTw. Embryos were then washed first at 3 μL/mL then 6 μL/mL of acetic anhydride/1% triethanolamine in PTw. These were washed out in PTw then fixed in 4% PFA for one hour on a rocker at room temperature. PFA was washed out with PTw and embryos were pre-washed in hybridization buffer (hyb) (50% formamide, 5× SSC, 50 μg/mL heparin, 0.1% Tween-20, 1% SDS, 100 μg/mL herring sperm DNA, 370 μL 1M citric acid) for 10 minutes at room temperature. This was replaced with a second hyb wash, and sealed vials were kept at 63°C for at least 12 hours.

Following pre-hybridization, DIG-labeled RNA probes were diluted 1:1000 in hybridization buffer and preheated to 90°C. Animals were transferred to mesh baskets in 24-well plates at 63°. All subsequent washes were performed by quickly transferring baskets to new wells with fresh solution. Probe was added to wells and baskets were transferred quickly into probe and left at 63°C for at least 48 hours. Probe was removed and stored for reuse, embryos were washed twice with hybridization buffer, and stepped 25%, 50%, and 75% SSC concentrations in hybridization buffer up to 100% 2x SSC solution, at 63°C. Next embryos were washed in 0.02x SSC, removed from heat and stepped into 25%, 50%, and 75% PTw in 0.02x SSC, up to 100% PTw. Embryos were then washed in 1x Roche blocking reagent for 1-2 hours at room temperature. Block was replaced with alkaline phosphatase-labeled (AP) anti-DIG Fab fragments (Roche) in Roche blocking reagent at a concentration of 1:5000, rocking overnight at 4°C. The next day antibody was removed, and embryos were washed 10 times for 15 minutes each in phosphate buffered saline with 0.1% Triton-X, then AP reaction buffer (100 mM NaCl, 50 mM MgCl_2_, 100 mM Tris, pH 9.5, 0.5% Tween-20) minus MgCl_2_, followed by two washes of AP buffer with MgCl_2_. Finally embryos were placed in BCIP-NBT (Promega) with AP buffer to react. Solution was checked for color change and replaced every half hour for the first several hours then once a day up to several weeks at 4°C, depending on speed of the reaction. When the chromogenic reaction was finished, embryos were washed in order with: PTw, water, ethanol, water, and PTw, then postfixed with PFA and cleared with 90% glycerol. Embryos were mounted on slides and imaged with DIC optics on a Zeiss Axioimager microscope.

## Additional Files

**Additional File 1.** Opsin alignment in Phylip format.

**Additional File 2.** Opsin gene tree file.

**Additional File 3.** Supplementary Tables 1-4.

Table S1. Opsin counts and recently published Anthozoan accessions

Table S2. Genomic and cross-reference evidence for *N. vectensis* opsins

Table S3. Opsin counts from each dataset from NvERTx

Table S4. Primers used for *in situ* hybridization

**Additional File 4.** Supplementary Figures S1-S3

Fig. S1. RNA-seq library analysis

Fig. S2. Opsin expression heatmaps from NvERTx datasets

Fig. S3. Sequence similarity of similar opsin genomic loci

## Declarations

Availability of data and materials

All RNA-seq raw reads generated during the current study are available in NCBI under BioProject ID PRJNA962884. Full length opsin sequences, transcriptome assembly, alignment and tree files are publicly available on DRYAD: https://datadryad.org/stash/share/qyUWZYiB-QKUMyLYuw0uMvLeOIkA-mAuNPU5_8NeeDQ. All other data are found in supplement included with this publication.

## Authors’ contributions

KJM, LSB, and KMK conceived of and designed the study. Funding acquisition by KMK and MQM. KJM, AL, LSB, and CD generated *in situ* probes and performed experiments. KJM and AL performed phylogenetics and sequence analysis. KJM wrote the manuscript. All authors read and provided feedback on the manuscript.

## Acknowledgements

The authors acknowledge the Minnesota Supercomputing Institute (MSI) at the University of Minnesota for providing resources that contributed to the research results reported within this paper. URL: http://www.msi.umn.edu. We’d like to thank Cassandra Extavour and Amaneet Lochab for help with microscope use.

