## Additional File 4 for "*Nematostella vectensis* exemplifies the exceptional expansion and diversity of opsins in the eyeless Hexacorallia"

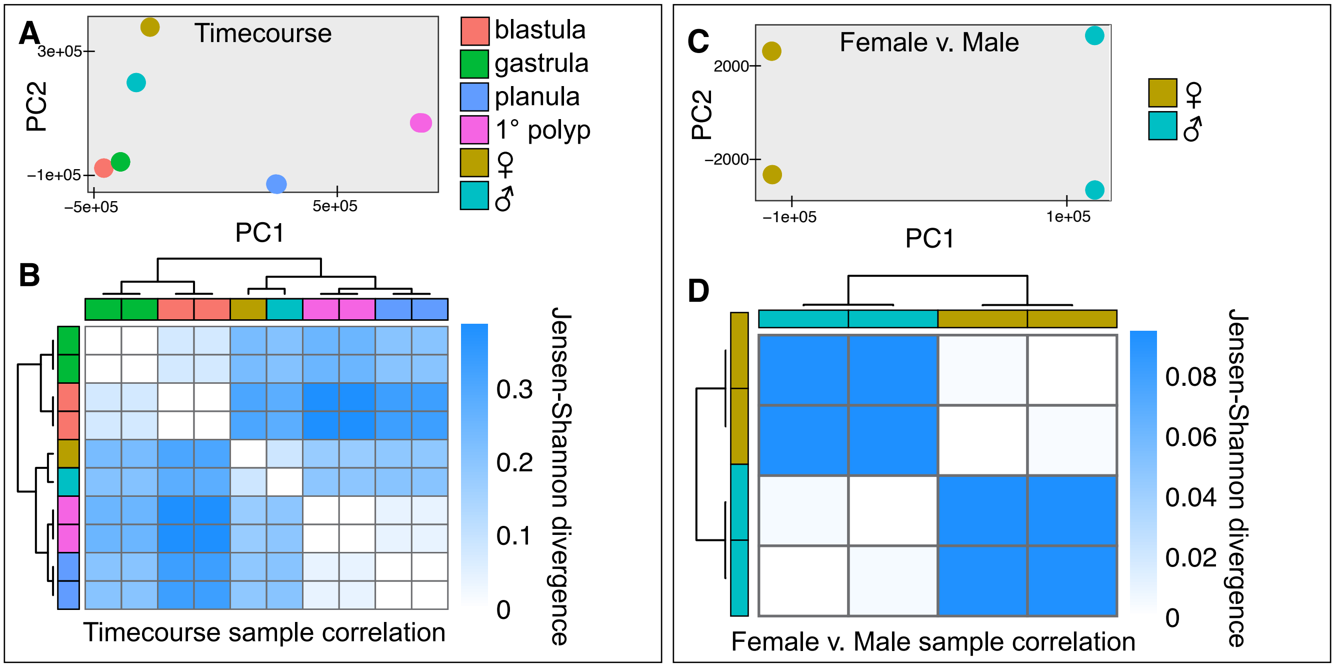


**Figure S1. RNA-seq library analysis.** **A)** Principle component analysis of RNA-seq timecourse libraries. Two libraries for each stage are shown, often directly over one another, indicating a high level of library similarity. One female and one male are shown. **B)** Jensen-Shannon divergence heatmap. More blue color signifies greater divergence. **C,D)** Same plots as A and B, except with for the two male and two female adults only.

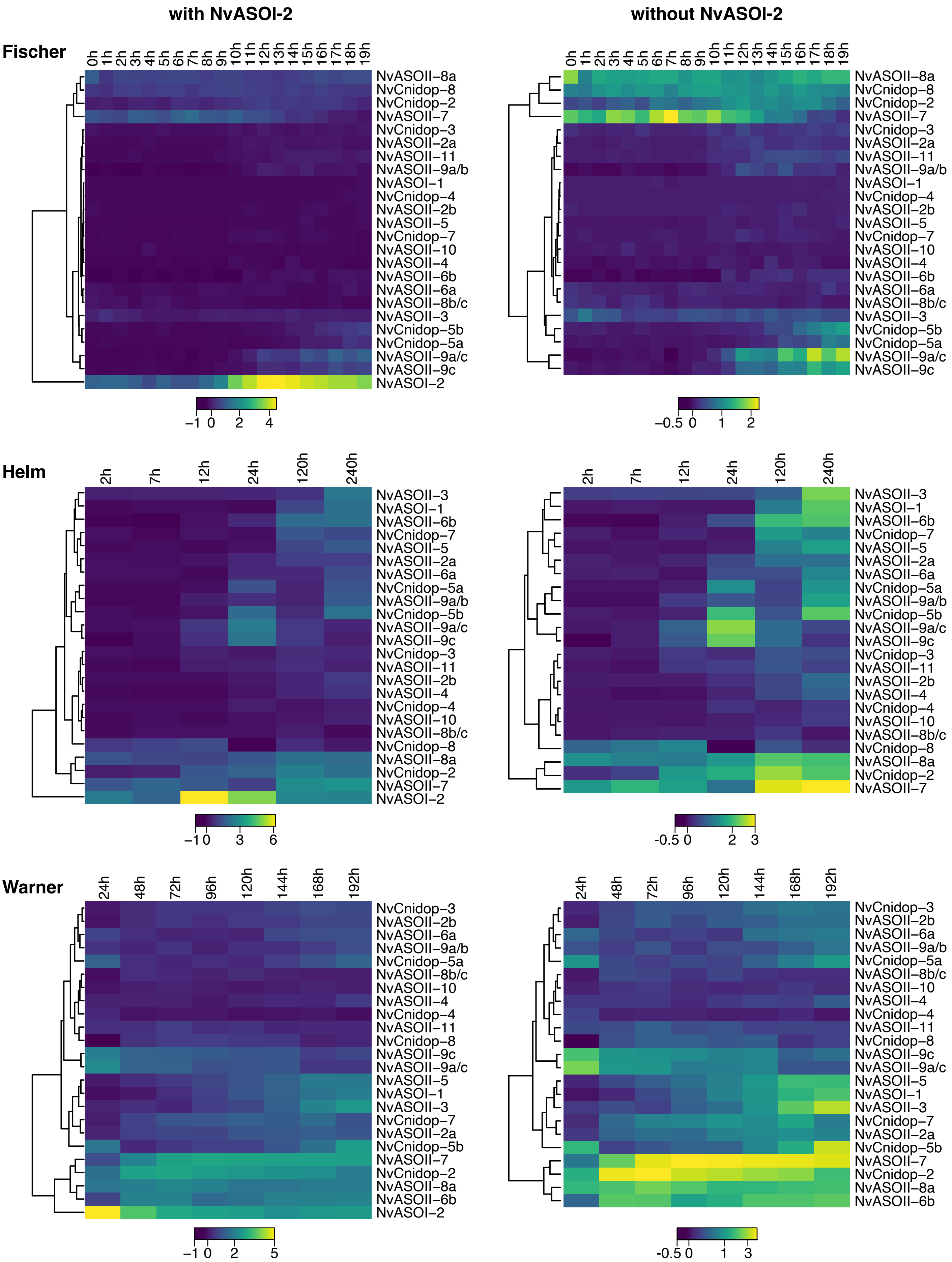

with NvASOI-2
without NvASOI-2
Fischer
Helm
Warner
0h 1h 2h 3h 4h 5h 6h 7h 8h 9h 10h 11h 12h 13h 14h 15h 16h 17h 18h 19h
2h 7h 12h 24h 120h 240h
24h 48h 72h 96h 120h 144h 168h 192h
NvASOII-8a
NvCnidop-8
NvCnidop-2
NvASOII-7
NvCnidop-3
NvASOII-2a
NvASOII-11
NvASOII-9a/b
NvASOI-1
NvCnidop-4
NvASOII-2b
NvASOII-5
NvCnidop-7
NvASOII-10
NvASOII-4
NvASOII-6b
NvASOII-6a
NvASOII-8b/c
NvASOII-3
NvCnidop-5b
NvCnidop-5a
NvASOII-9a/c
NvASOII-9c
NvASOI-2
−1 0 2 4
−0.5 0 1 2
−1 0 3 6
−0.5 0 2 3
−1 0 2 5
-0.5 0 1 3

**Figure S2. Opsin expression heatmaps from NvERTx datasets.** Heatmaps are derived by embryonic count data listed for transcripts from each of three databases on NvERTx (Helm et al. 2013, Fischer et al. 2014, Warner et al. 2017). Names on each row correspond to each database. Left column displays heatmap with the highly expressed *NvASOI-2,* while right column displays heatmaps without this transcript included. BLAST was used with each final paralog from this study as bait on NvERTx, and the top hits from NvERTx were aligned to our paralogs and manually checked for highest sequence identity. Some hits from similar but distinct paralogs were combined into the same count data on NvERTx, thus some overlapping sequences are found in the same row for these heatmaps. For reference between our paralog IDs, sequence, and NvERTx count see Table S2 and Additional File S4.

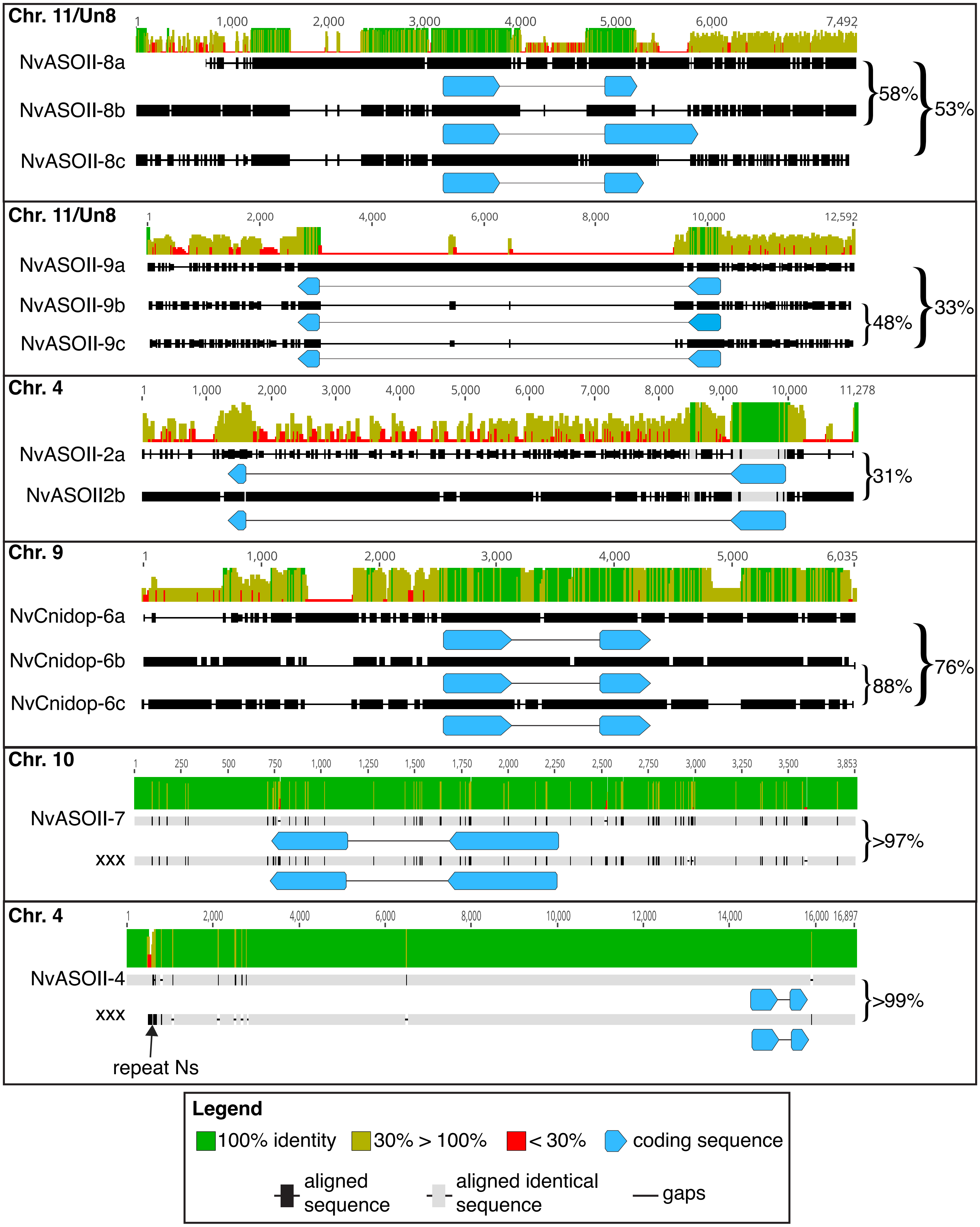


**Figure S3. Sequence similarity of similar opsin genomic loci.** The genomic loci for opsins with highly similar coding sequences are shown in each box. Gene names and chromosome(s) on which they are found are listed for each comparison (for coordinates, see Table S2). Scale is in base pairs including gaps. Each sequence is represented by the black lines (thin line, gaps; thick black, non-identical alignment; thick gray, identical alignment). Consensus identity is represented by color and height of bars at the top of each comparison and pairwise identity of the most similar pair, and consensus identity are listed to the right. Similar sequences were not found in BLAST results of the Wellcome-Sanger genome.
